# Functional Reconstitution of Human OATP1B1 and OATP1B3 in Proteoliposomes: Effect of Bicarbonate Gradients and Substrate-specific Stimulation on Transport

**DOI:** 10.64898/2026.10.29.762842

**Authors:** Juan Carlos Mellado Tenorio, Stephen J. Butler, Bruno Stieger, Kaspar P. Locher

## Abstract

The hepatic organic anion transporting polypeptides OATP1B1 and OATP1B3 are critical for the clearance of widely prescribed drugs, hepatotoxins, and endogenous metabolites. However, the thermodynamic driving force governing their transport has remained a subject of debate due to the complexity of cellular models. Here, we report a defined, reconstituted proteoliposome system to study the transport kinetics of isolated human OATP1B1 and OATP1B3. Through direct functional assays, real-time counter-ion tracking, and a novel ABCG2-OATP1B1 functional co-reconstitution, we identify an outwardly directed bicarbonate gradient as a thermodynamic driving force for concentrative substrate uptake. In contrast we find no evidence for the proposed role of reduced glutathione efflux as a possible driving force. Targeted mutagenesis experiments revealed that the conserved TM3 residue His115 is needed to couple the bicarbonate gradient to concentrative substrate uptake, mechanistically linking substrate antiport to local pH-sensitive bicarbonate processing. We also investigated the substrate-specific stimulatory effect of the green tea catechin epigallocatechin gallate (EGCG) on OATP1B3 transport, confirming that this effect is intrinsic to OATP1B3, substrate specific, and does not need the involvement of other cellular proteins. Our findings define the energetic basis of OATP1B1 and OATP1B3 hepatic xenobiotic clearance.

## INTRODUCTION

The liver plays a vital role in the clearance of endogenous compounds and xenobiotics. The rate at which the latter—whether therapeutics or toxic compounds—are cleared from the bloodstream by the liver depends on two factors: hepatocyte uptake and subsequent enzymatic metabolism (e.g., via cytochrome P450s and/or conjugation). Organic anion transporting polypeptide (OATP) 1B1 and OATP1B3 (SLCO1B1 and SLCO1B3) are key players in the uptake across the basolateral membrane of the hepatocytes, mediating portal-to-liver uptake of a vast array of molecules (*1–3*). Their clinical significance is profound: the EMA and FDA include OATP1B1 and OATP1B3 in their core set of transporters to be evaluated during preclinical drug development, mandating strict labelling for known inhibitors. Common substrates include widely prescribed drugs such as statins, angiotensin receptor blockers (e.g., valsartan), antibiotics (e.g., rifampicin), and antivirals (e.g., ritonavir) (*4, 5*). Furthermore, OATP1B1 and OATP1B3 are the primary entry routes for the strongly hepatotoxic α-amanitin from the Amanita phalloides mushroom, the clinical treatment of which relies on potent OATP inhibitors to block liver uptake (*6*).

Given their pharmacological importance, OATP1B1 and OATP1B3 uptake assays are an indispensable step in drug development. Consequently, the kinetics, regulation, genetics, and transport mechanisms of these proteins have been extensively studied (*1, 7–12*). Historically, these investigations have relied on diverse cellular systems, including primary hepatocytes and overexpression or knockout models in CHO and HEK293 cells. While these models have provided important insights into drug-drug interactions, they frequently yield conflicting data. The discrepancies are largely due to the inherent complexities of the cellular environment, such as variable endogenous protein expression and competing metabolic pathways. To date, a purely reconstituted protein system for OATPs has not been reported. Establishing such a system is critical, as it would provide a uniform, unambiguous model for affinity assays, effectively disentangling the act of transport from downstream metabolic disposition as well as investigation of the impact of membrane lipid composition on OATP function (*13*).

This methodological shift from cellular models to isolated, reconstituted environments has strong historical precedence across the Solute Carrier (SLC) superfamily, particularly for proteins governed by ion coupling or obligate exchange. A classic early example is the Excitatory Amino Acid Transporter family (SLC1/EAATs). By isolating the carrier in reconstituted proteoliposomes researchers were able to systematically control both sides of the bilayer and definitively split the transport cycle into its distinct sodium-dependent influx and potassium-dependent reset phases (*14, 15*). Beyond amino acid carriers, resolving the energetic landscape of anion exchangers has faced hurdles. The exact stoichiometry and electrogenicity of the chloride/bicarbonate exchanger SLC26A6 remained a long-standing debate due to the overlapping anion channels and continuous intracellular metabolic buffering of cells. Only by reconstituting the purified transporter into proteoliposomes could researchers unambiguously establish its coupled transport mechanism (*16*). Similarly, the biophysical nature of CLC proteins was redefined when a CLC homologue was reconstituted into the leak-free environment necessary to impose strict, isolated pH and chloride gradients. While long assumed to be chloride channels, this was vital to unmasking a strict 2Cl^−^ : 1H^+^ stoichiometric coupling, proving that these proteins operate as secondary active antiporters—a discovery that native proton leaks in living cells would have otherwise masked (*17*). Finally, precise stoichiometric debates are routinely settled *in vitro*, as demonstrated by the Na^+^/Ca^2+^ exchanger (NCX). By utilising a proteoliposome system, researchers were able to perfectly clamp the membrane potential alongside precise internal and external ion concentrations, definitively proving an exchange stoichiometry (*18*). These precedents demonstrate that for transporters governed by delicate counter-ion loops, a purified platform is an absolute prerequisite.

Despite recent structural insight, the exact thermodynamic driving force of OATP1B1 and OATP1B3 has remained a subject of debate (**Fig. 1A**). Early studies utilising rat hepatocytes established that alkali-loaded cells exhibited increased taurocholate uptake, with transport helping to recover intracellular pH. This suggested an organic anion/ OH^-^(or HCO_3_^-^) exchange mechanism (*19*). Subsequent work linked this pH dependence to a conserved histidine residue on the extracellular side of transmembrane domain 3 (TM3), demonstrating that OATPs lacking this histidine lose their pH sensitivity. This led to a proposed HCO_3_^-^ extrusion model, sustained by: (I) a basolateral Na^+^/HCO_3_^-^ symporter; (II) the acidic microenvironment within the Space of Disse (perisinusoidal space), maintained by the Na^+^/H^+^ exchanger, which would facilitate the conversion of HCO_3_^-^ exported to CO_2_ (*20*); and (III) the presence of carbonic anhydrase in the hepatocyte membrane which would catalyse that conversion(*21*). Other studies utilising electrophysiology suggested that while rat OATP1A1 functions as an electroneutral exchanger, OATP1B1 and OATP1B3 may operate as electrogenic transporters (*22*). Further complicating the landscape, intracellular reduced glutathione (GSH) has been proposed as an alternative driving force, although conflicting evidence was found between rodent and human isoforms (*23, 24*). Recently, high-resolution structural studies have provided a critical clue: a triangular density interacting with His115 in the apo state of OATP1B3, interpreted as a bound HCO_3_^-^ion (*10*). However, a limitation of the existing literature is that functional evidence on the role of bicarbonate has relied on the measurement of transport-modulating pH alkalinisation or acidification, rather than direct tracking of bicarbonate translocation. Furthermore, it has remained highly challenging to distinguish the translocation of a weak base like HCO_3_^-^ from counter-transport of H^+^ or the co-transport of OH^-^ as these result yield similar results in bulk pH assays. Consequently, definitive functional proof of the identity of the co-transported ion has remained elusive.

**Figure 1.**
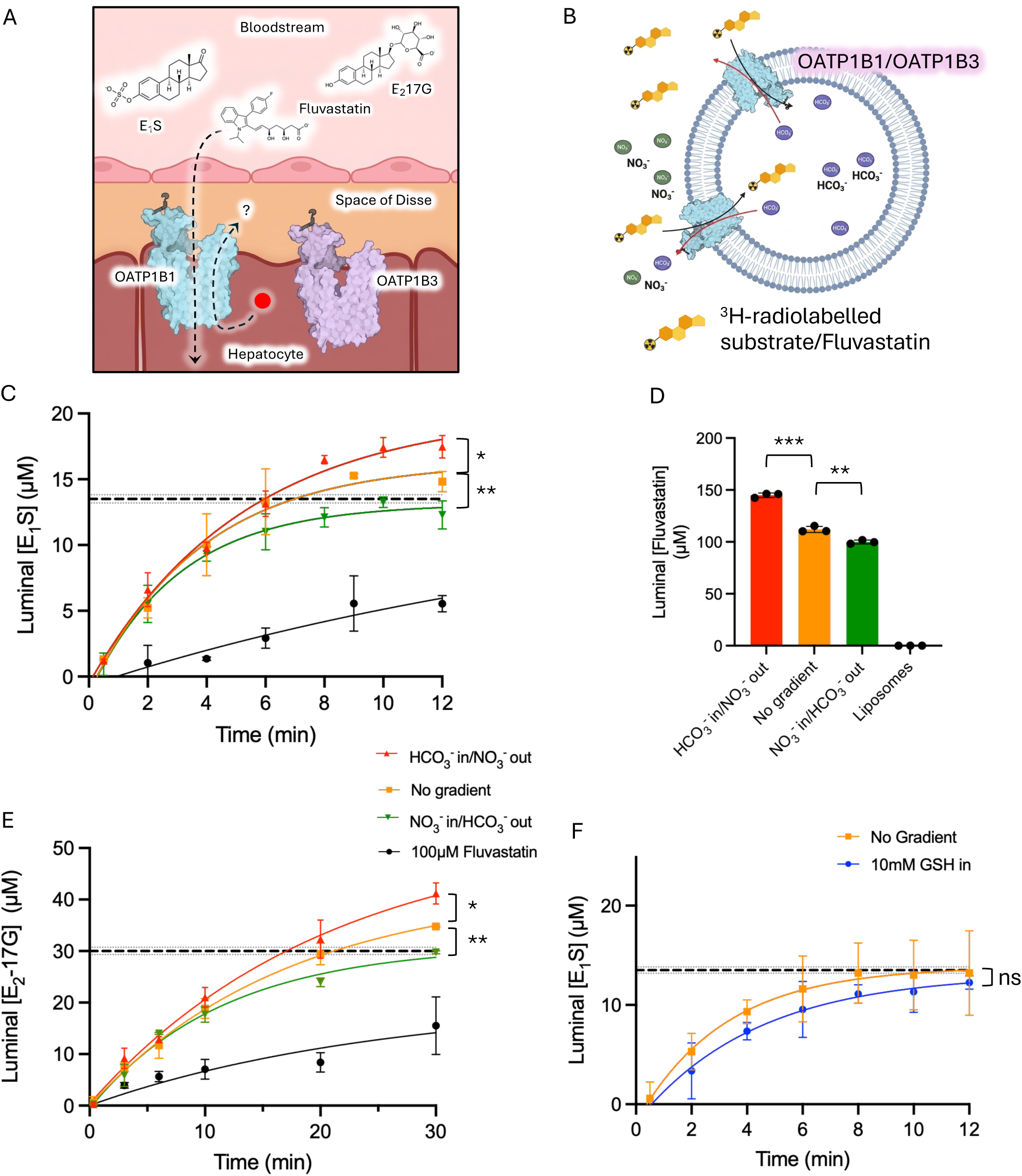
Bicarbonate gradients drive OATP1B1 and OATP1B3-mediated transport in proteoliposomes. **A:** Schematic of OATP1B1-and OATP1B3-mediated uptake of endogenous compounds and xenobiotics into the hepatocyte. The theoretical driving force or counterion is depicted with a red circle. **B:** Schematic of the experimental setup to test the effect of opposing HCO_3_^−^ gradients on OATP1B-mediated transport. Proteoliposomes were loaded with either 15mM NaHCO_3_ or 15mM NaNO_3_ and diluted into external buffers containing the reciprocal salt to establish an outwardly-directed bicarbonate gradient (*[HCO_3_ ^−^*]*_out_>[HCO_3_ ^−^]_in_*) or an inwardly directed gradient (*[HCO_3_ ^−^*]*_in_>[HCO_3_ ^−^]_out_*) respectively, while maintaining osmotic equilibrium. **C:** Time-course of ^3^H-E_1_S uptake by OATP1B1. Transport is significantly enhanced by an outward HCO_3_^−^ gradient compared to inward gradient or symmetrical (no gradient) conditions. **D:** Accumulation of the statin fluvastatin in OATP1B1-reconstituted proteoliposomes under the indicated HCO_3_^−^ gradient conditions. **E**: OATP1B3-mediated uptake of ^3^H-E -17G into proteoliposomes driven by an outward HCO_3_^−^ gradient. **F**: Effect of an outwardly-directed gradient of 10mM GSH on OATP1B1-medited ^3^H-E_1_S uptake. To dissipate the electrochemical gradient and maintain ionic balance, 5 mM potassium tartrate was supplemented in the external buffer. Data represent mean ± s.e.m. of 3 independent experiments. ns: *p* > 0.05, *: *p* < 0.05, **: *p* < 0.01, ***: *p* < 0.001.

Another unresolved mechanistic question concerns how OATP1B1 and OATP1B3 exhibit substrate-specific stimulation or inhibition in response to green tea catechins, a phenomenon with potentially important pharmacokinetic consequences (*25–28*). Although this unusual behaviour has been studied extensively in cellular systems, it has not yet been examined at the level of the purified proteins.

In this study, we sought to investigate the driving force and transport mechanism of OATP1B1 and OATP1B3 by stripping away additional variables of cellular systems, as well as to look at the substrate specific effect of the modulator green tea catechin EGCG on OATP1B3-mediated transport. By successfully reconstituting purified human OATP1B proteins into proteoliposomes, we established a well-defined, *in vitro* system to isolate their kinetics. Through mechanistic studies, we demonstrated that OATP1B1 and OATP1B3 can act as bicarbonate-driven exchangers, where an opposing HCO_3_^-^ gradient provides the thermodynamic power for concentrative substrate uptake. Furthermore, through targeted mutagenesis of the conserved His115 residue, we propose a kinetic model wherein these transporters utilise a HCO_3_^-^ catalytic mechanism to drive the efficient clearance of diverse xenobiotics.

## RESULTS

### Functional reconstitution of OATP1B1 and OATP1B3 in proteoliposomes

Multidrug transporters, and specifically polyspecific solute carriers, present particular challenges for functional reconstitution: their intrinsic instability due to the large conformational changes needed to transport bulky, chemically diverse substrates; their strong dependence on specific membrane lipid interactions and the amphipathic nature of the substrates complicate the development of reproducible, proteoliposome-based transport assays. OATP1B1 or OATP1B3 are highly dynamic and labile membrane proteins. In consequence, iterative optimisation of numerous steps in the reconstitution process was required, including the identification of a suitable mixture of synthetic and natural lipids (see **Methods** section), and the optimal Triton X-100 concentration for liposome destabilisation (**Fig. S1A**). This approach yielded a robust experimental procedure with consistent reconstitution efficiency (**Fig. S1B**) and the compartmentalisation necessary to establish asymmetric ionic conditions across the lipidic membrane (**Fig. 1B**). The chemical nature of OATP1B1 and OATP1B3 substrates, which combine hydrophobic and anionic properties, also required the optimisation of the filter material used for transport assays. We found that PVDF filters with a 0.2µm pore size provided the lowest amount of unspecific substrate binding if they were rinsed with ice-cold buffer with high concentrations of unlabelled substrate both before and after filtration. This system allowed us to evaluate the coupling of substrate flux to trans-membrane ion gradients.

Utilising the calcein entrapping method (*2S, 30*), the internal aqueous volumes of these liposomes after extrusion through 0.4µm filters were measured to be 0.85 ± 0.02 µL per mg of lipids (**Fig. S1C**). Accurately measuring the entrapped volume allows reliable determination of luminal substrate concentrations, which is particularly important for studying low turnover transporters such as the OATPs, and to differentiate active, concentrative transport from passive facilitated diffusion. This experimentally measured luminal volume allowed us to determine the intra-vesicular substrate accumulation required to reach substrate equimolarity with the external buffer, which is depicted by the horizontal dotted lines in our transport assay plots.

### An outwardly directed HCO_3_^-^ gradient can power OATP1B1 and OATP1B3-mediated transport

We first showed that, in this proteoliposome system, OATP1B1 could transport ^3^H-Estrone-3-sulfate (^3^H-E_1_S) into proteoliposomes in a mildly concentrative manner, reaching around 1.1-fold substrate concentration in the lumen compared to the outside of the proteoliposomes, and this transport could be competitively inhibited by fluvastatin (**Fig. 1C**). We called this condition “no gradient” since no bicarbonate gradient was imposed. However, this cannot be considered a “bicarbonate-free” condition, as continuous exchange from atmospheric CO_2_ supplies trace amounts of HCO_3_^-^, which we estimate to be around 0.2mM (assuming atmospheric *p*CO_2_=400 ppm, and using Henry’s law and a buffer pH of 7.4). In a separate experiment the proteoliposomes were then loaded with 15mM NaHCO_3_, while 15mM NaNO_3_ was added to the outside buffer to equalise for osmolarity and ionic strength of the external and luminal compartments. This outwardly directed HCO_3_^-^ gradient enhanced ^3^H-E_1_S transport; while we could not accurately determine the initial velocities, the concentrative power of the transporter increased to 1.3-fold the concentration of substrate in the external medium. Interestingly, when the ion gradient was inverted, transport was reduced to an equilibrative reaction. The steady state concentrations were found to be significantly different (extra sum-of-squares F tests, *p* = 0.0069, F = 8.32 for “HCO_3_^-^out” vs. “no gradient”, *p* =0.0158, F = 6.477 for “no gradient” vs “HCO_3_^-^ in”). Moreover, E_1_S accumulation correlated with the magnitude of the bicarbonate gradient, and the response plateaued between 15-20mM HCO_3_^-^ (Fig. S2B). Importantly, nonspecific adsorption of E_1_S to the hydrophobic liposome membrane was detected in protein-free proteoliposomes (Fig. S2A), and this signal was subtracted to that of OATP1B1-conatining liposomes for an accurate determination of the luminal substrate concentration. These results constitute the first evidence, in a purified protein system, that OATP1B1 can use an outwardly directed HCO_3_^-^ gradient to support concentrative substrate import.

To validate this mechanism with a physiologically relevant and chemically distinct substrate, we examined the effect of HCO_3_^-^ on the transport of fluvastatin. We used reverse-phase ultra-high pressure chromatography and the intrinsic absorbance of statins at 240 nm to integrate the elution peaks and quantify luminal concentrations for this substrate. Unlike E_1_S, hydrophilic statins such as fluvastatin exhibit minimal non-specific partitioning into the lipidic bilayer, which allows for a more precise quantification of the internal aqueous concentration. In the presence of an inhibitory (inwardly-directed) bicarbonate gradient, OATP1B1 mediated transport reached near equimolar conditions (99.9 ± 1.8 μM in the lumen vs. 100 μM external) (**Fig. 1D**). In contrast, a favourable outward HCO_3_^-^gradient drove fluvastatin accumulation to approximately 1.4-fold the external concentration (145 ± 2 µM). Notably, in the absence of a bicarbonate gradient or any added bicarbonate, OATP1B1 still achieved a modest concentrative effect (112 ± 3 µM), which may be attributed to trace amounts of bicarbonate present in the reaction buffer. These values were found to be significantly different (unpaired t-test, *p* = 0.0002, F = 1.84 for “HCO_3_^-^ out” vs. “no gradient”, *p* =0.072, F = 2.8513 for “no gradient” vs “HCO_3_^-^ in”), and closely match the magnitudes achieved in the ^3^H-E_1_S transport assays. Collectively, this data demonstrate that bicarbonate plays a fundamental role in the transport mechanism of OATP1B1, as the directionality of the bicarbonate gradient affects the uptake of both the model substrate E_1_S and the clinically used drug fluvastatin.

We then sought to validate whether this mechanism is conserved in OATP1B3. Using a similar proteoliposome setup, we measured the accumulation of [^3^H]-estradiol-17-β-D-glucuronide ([^3^H]-E_2_17G) over time. In the case of OATP1B3, this substrate is preferred over E_1_S due to its lower K_M_ for the transporter (5-25 µM for E_2_17G and too low for accurate determination in the case of E_1_S) (*1, 4*).

In our setup, E_2_17G also shows less nonspecific adsorption, akin to fluvastatin (Fig. S2C). We found that an inwardly directed HCO_3_^-^ gradient resulted in an equilibrative [^3^H]-E_2_17G transport process, whereas an outward gradient led to an approximately 1.4-fold concentration of substrate in the lumen. In the absence of a gradient, a modest approximate 1.15-fold concentration of [^3^H]-E_2_17G was reached (**Fig.1E**). Statistical analysis also found the steady-state substrate concentrations to be significantly different (extra sum-of-squares F tests, *p* = 0.0158, F = 6.477 for “HCO_3_^-^ out” vs. “no gradient”, *p* =0.0007, F = 14.90 for “no gradient” vs “HCO_3_^-^ in”). Strikingly, these findings closely mirror the values obtained for OATP1B1 experiments and confirm the role of bicarbonate as a likely driving force OATP1B1 and OATP1B3-mediated transport, where an outwardly directed gradient facilitates concentrative substrate uptake.

Finally, we investigated the proposed role of reduced glutathione (GSH) as a driving force. We loaded proteoliposomes with 10mM GSH and resuspended them in a buffer containing 5mM potassium tartrate, to maintain charge neutrality and prevent the development of a membrane potential. The resulting transport reaction largely overlapped with the control obtained in the absence of a gradient (**Fig.1F**). Comparison of the non-linear regression models using an extra sum-of-squares F-test confirmed that a single global fit sufficiently described both datasets, with no significant differences observed in either the transport rate (*k*) or the steady-state plateau (p=0.4876, F=0.4919). Thus, we found no evidence that a GSH gradient affects transport in our experimental system.

### Co-reconstitution with ABCG2 confirms substrate/HCO_3_^-^antiport by OATP1B1

While our initial results demonstrated that a bicarbonate gradient can energise OATP1B-mediated transport, we sought to further investigate the role HCO_3_^-^as a counter-ion in the transport cycle. To achieve this, we co-reconstituted purified OATP1B1 and the ATP-binding cassette (ABC) transporter ABCG2 (**Fig. 2A**). A molar ratio of 2 OATP1B1 molecules per ABCG2 dimer provided the best results in our assays, and several freeze thaw cycles were carried out prior to proteoliposome extrusion to ensure an even, randomised distribution of both OATP1B1 and ABCG2 across all membranes. By leveraging their common substrate specificity for E_1_S, we created a vectorial relay: the primary active transporter ABCG2 serves as a pump to drive unidirectional substrate import (as ATP is only present only in the external buffer, which allows concentrating E_1_S in the lumen), while OATP1B1 can import or export substrate depending on the prevailing electrochemical gradients, and therefore will export substrate once a sufficient internal concentration is reached. With this coupled assay, we could create a controlled luminal environment in real time and measure the transport activity of OATP1B1 in response to a changing substrate concentration gradient. Addition of ATP initiated the accumulation of [^3^H]-E_1_S in the proteoliposome lumen. This accumulation reached a steady-state plateau after approximately 30 min, representing the kinetic equilibrium between ABCG2-mediated import and OATP1B1 mediated export (**Fig. 2B**). To confirm that OATP1B1 was responsible for the export flux, we employed atazanavir, a selective inhibitor for OATP1B1 over ABCG2 (IC_50_1.4 µM vs. IC_50_ 69.1 µM, respectively) in the right concentration range (*31, 32*). Inhibition of OATP1B1 resulted in a sustained 40% increase in [^3^H]-E_1_S accumulation over 60 min, with no signs of plateauing in the measured time frame. Crucially, the addition of 15 mM NaHCO_3_ to the external buffer shifted the steady-state toward a lower luminal concentration, indicating the presence of a bicarbonate gradient stimulated OATP1B1-mediated export. This further confirms that the HCO_3_^-^ gradient modulates the transport rate of OATP1B1. We then validated this result by the addition of the ABCG2 inhibitor Ko143 at the 29-minute mark, which triggered an immediate net loss of substrate, as OATP1B1-mediated export was no longer offset by ABCG2 activity.

**Figure 2.**
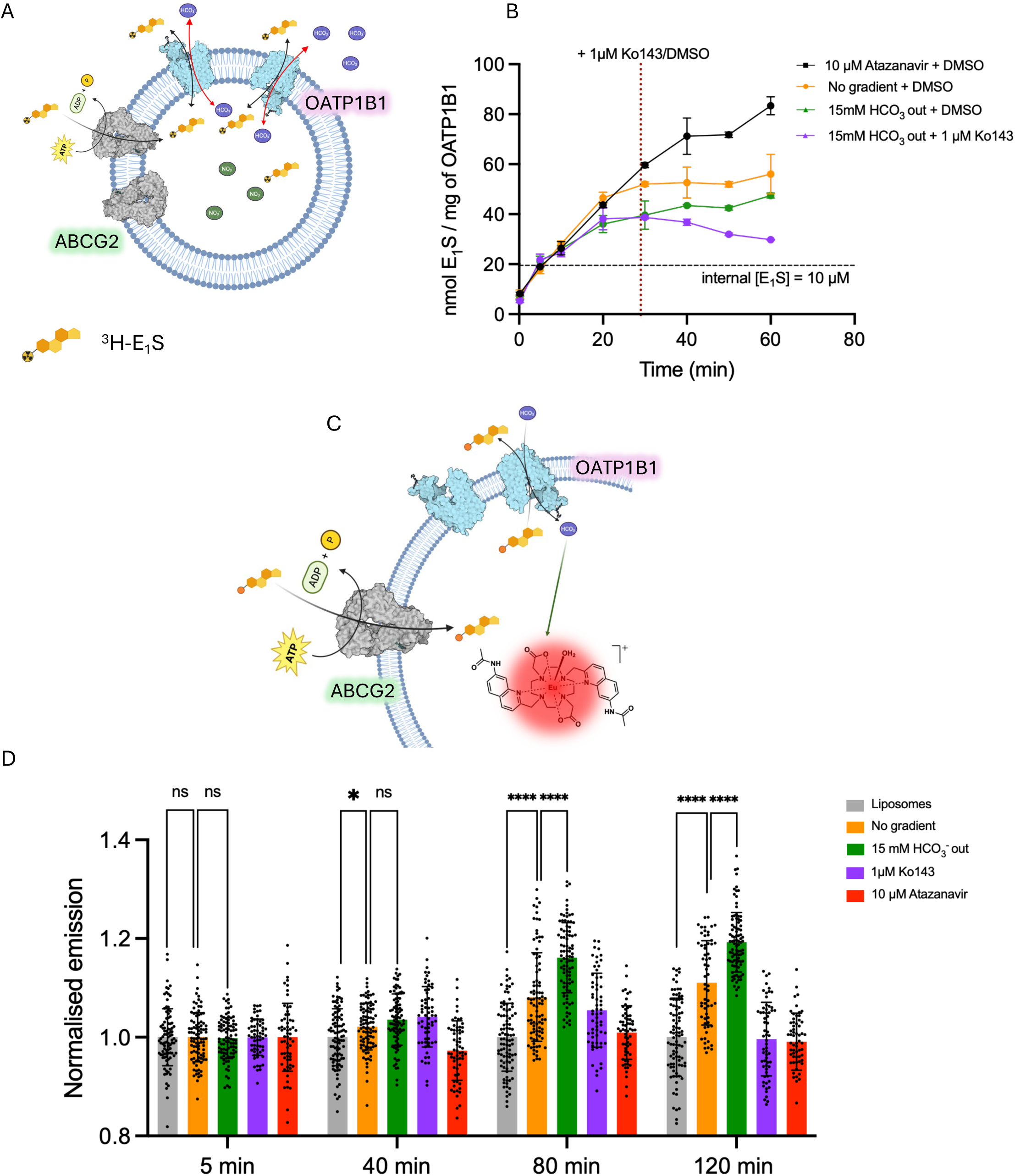
OATP1B1-mediated transport is coupled to bicarbonate antiport in poreoliposmes containing OATP1B1 and ABCG2. **A:** Schematic of the co-reconstitution system in proteoliposomes. OATP1B1 is co-inserted with the ABC transporter ABCG2. The proteoliposomes were preloaded with 15mM NaNO_3_ if 15mM NaHCO_3_ was added to the outside buffer (HBS pH 7.4). Transport was initiated by the addition of ^3^H-E_1_S and ATP. **B:** Time-course of ^3^H-E_1_S accumulation into the proteoliposomes. 1 µM Ko143 or DMSO was added at t = 29 min. Horizontal dotted lines represent the theoretical equilibrium (equimolar internal and external E_1_S concentrations). **C:** Schematic of the HCO_3_^-^detection system in the co-reconstituted liposomes. **D:** Real-time monitoring of luminal HCO_3_^-^flux using the Eu(III) luminescent probe. Normalised emission traces show the change in internal HCO_3_^-^concentration during OATP1B1-mediated subtrate exchange. Data represent mean ± s.e.m. of 3 independent experiments. ns: *p* > 0.05, *: *p* < 0.05, **: *p* < 0.01, ***: *p* < 0.001.

### Direct, real-time monitoring of HCO_3_^-^flux

To provide direct evidence of bicarbonate transport, we employed a specialised lanthanide-based luminescent probe, the Eu(III) probe [Eu.L^1^]^+^ which allowed us to monitor luminal HCO_3_^-^in real-time. This probe was originally developed to bind and detect fluoride, but it was found to be also responsive to bicarbonate and was subsequently adapted into a liposome-based assay for monitoring this anion (*33, 34*). [Eu.L^1^]^+^ exhibits high selectivity for bicarbonate over chloride and nitrate, with negligible changes in emission intensity or spectral shape observed for the latter anions even at 1000-fold excess, whereas bicarbonate binding produces a pronounced increase in emission intensity and lifetime (*32*). Furthermore, its application to monitoring bicarbonate transport into liposomes has been validated in HEPES buffer containing high concentrations of nitrate and chloride (*33, 34*), making it an ideal probe for the present study.

A significant technical challenge was the sensitivity mismatch between the low micromolar concentration of [^3^H]-E_1_S and the millimolar detection thresholds of the HCO₃⁻ probe. To overcome this, we utilised our co-reconstitution system to amplify the signal: ABCG2 continuously resupplied E_1_S into the lumen, enabling a prolonged OATP1B1-mediated export, and a concomitant HCO_3_^-^accumulation time. Proteoliposomes were pre-loaded with the Eu(III) probe and 100 µM E_1_S, and subsequently resuspended in a buffer containing a lower (50 µM) E_1_S concentration and an ATP-regeneration system. This experimental design established an immediate, outwardly directed substrate gradient ([E_1_S]_in_ > E_1_S]_out_), ensuring that OATP1B1-mediated export—and the corresponding HCO_3_^-^import—commenced instantly. To sustain the transport reaction across longer time frames and avoid ATP depletion, we also employed a creatine phosphokinase-based ATP regeneration system.

Upon initiation of the reaction, we observed a progressive increase in luminescence (at 617 nm), signalling the import of HCO_3_^-^into the lumen (**Fig. 2D**). The luminescence increase was significantly more pronounced 80 min after 15 mM HCO_3_^-^was added to the external buffer, but abolished by the addition of atazanavir, confirming that the bicarbonate flux is OATP1B1-dependent. Furthermore, the addition of the ABCG2 inhibitor Ko143 resulted in only a transient increase in luminescence that rapidly plateaued; without the continuous replenishment of luminal E_1_S by ABCG2, the chemical gradient driving the exchange was quickly exhausted (**Fig. 2D**).

Collectively, these experiments demonstrate that OATP1B1-mediated transport is mechanistically coupled to the antiport of bicarbonate. Our results suggest that the gradient is a primary determinant of the energetic landscape and directionality of OATP1B-mediated transport.

### Mutagenesis of the putative bicarbonate binding site provides insight into the transport cycle

The recently reported structure of OATP1B3 featured a triangular-shaped density feature that was assigned to a bicarbonate ion, located in a pocket near the signature motif, and interacting with Histidine 155 (*10*). Based on structural evidence and previous studies (*20, 35*), we hypothesised that the homologous residue in OATP1B1, also His115, plays a crucial role in bicarbonate coordination (**Fig. 3A**). To test this, a H115A variant of OATP1B1 was purified and reconstituted into proteoliposomes, and its activity was measured under various bicarbonate gradients. We found that the directionality of the HCO_3_^-^gradient no longer affected ^3^H-E_1_S transport by OATP1B1_H115A_ (**Fig. 3B**). Furthermore, the mutation abolished the concentrative power of the transporter compared to the wild-type (WT) (**Fig. 3D**). However, the mutation did not markedly reduce transport rate: the variant still dissipated the substrate electrochemical gradient on a comparable timescale. These findings identify His115 as critical for bicarbonate-dependent coupling rather than for substrate translocation itself. Although the data do not establish that bicarbonate countertransport is completely abolished, they show that the energetic advantage conferred by the HCO_3_^-^gradient is lost in the mutant. This supports the conclusion that coupling substrate transport to HCO_3_^-^antiport is required for the concentrative capacity of OATP1B1, with His115 serving as a key residue in that mechanism.

**Figure 3.**
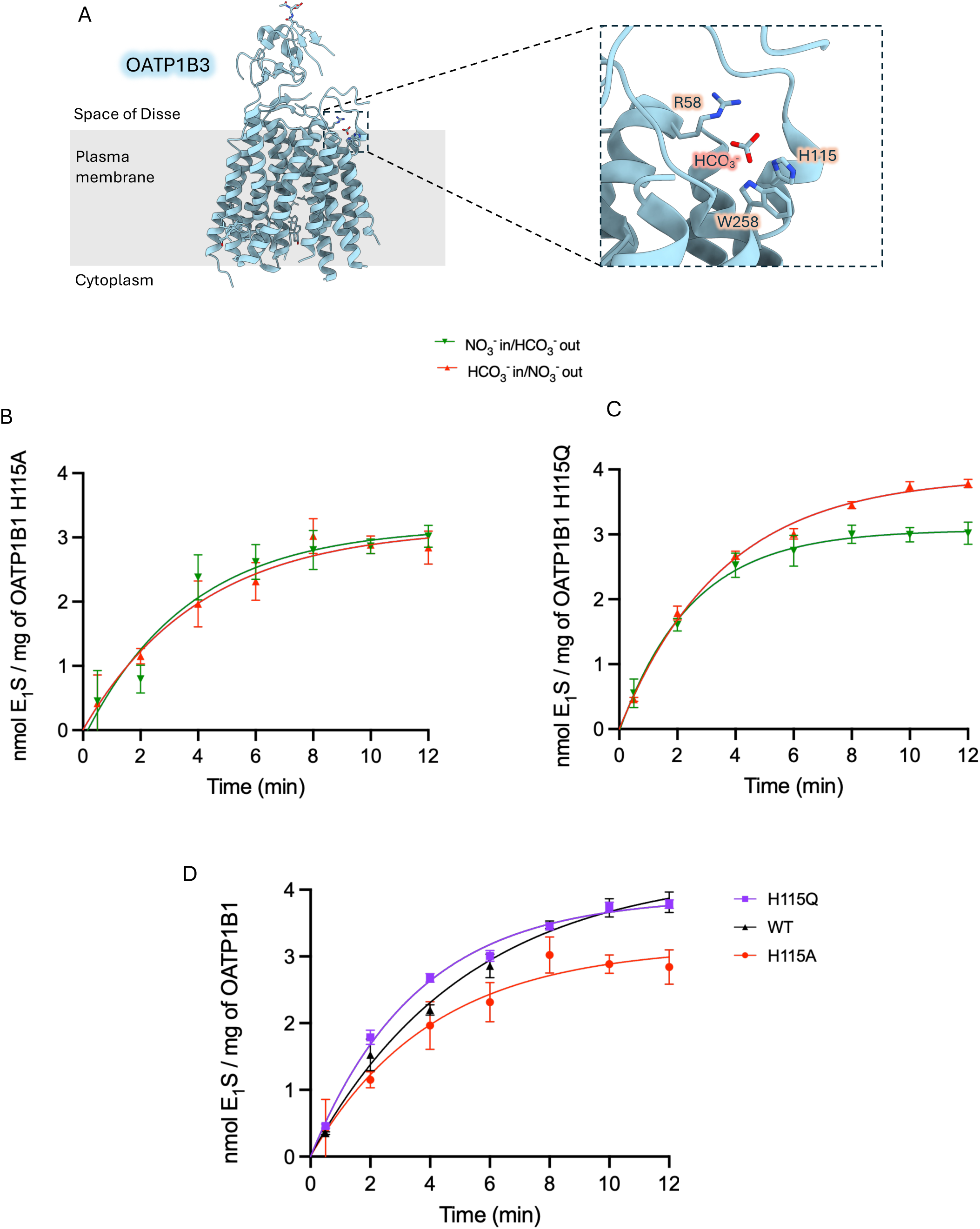
Mutations in the HCO_3_^-^binding site alter OATP1B1 transport activity. **A:** Close-up view of the bicarbonate binding site in OATP1B3 in its bicarbonate-bound conformation (PDB: 8PG0). **B:** OATP1B1 H115A-mediated uptake of ^3^H-E_1_S in reposnse to an outward or inward HCO_3_^-^gradient. **C:** OATP1B1 H115Q-mediated uptake of ^3^H-E_1_S in reposnse to an outward or inward HCO_3_^-^gradient. **D:** Uptake comparison between OATP1B1, H115A, and H115Q. For the three curves, an outwardly directed 15mM HCO_3_^-^gradient was used. Data represent mean ± s.e.m. of 3 independent experiments. Epigallocatechin gallate (EGCG)

To further probe the chemical requirements of this site, we introduced a H115Q mutation. We hypothesised that the glutamine side chain, as a hydrogen-bond donor, might partially mimic the coordination provided by histidine. Upon purification and reconstitution, we observed that the H115Q variant regained its sensitivity to the bicarbonate gradient (**Fig. 3C**) and successfully recovered its concentrative power (**Fig. 3D**). Interestingly, H115Q exhibited faster initial transport rates than the WT in this experimental system. While the mechanistic basis for this is explored further in the Discussion, this acceleration likely reflects a lower-affinity interaction with HCO_3_^-^that facilitates faster dissociation and subsequent conformational cycling.

### EGCG selectively stimulates the uptake of E_1_S by OATP1B3

Interactions between green tea flavonoids and the OATPs have been widely reported, with some compounds exhibiting complex, substrate-dependent modulatory effects on transport activity. Notably, the flavonoid epigallocatechin gallate (EGCG) was shown to stimulate the E_1_S transport rate by OATP1B3 in whole-cell assays (CHO cells), purportedly by increasing substrate affinity. Conversely, the same study reported that EGCG non-competitively inhibited the uptake of other substrates while having no measurable effect on the transport of E_2_17G (*26*). We sought to verify whether this observation can be reproduced in a chemically defined proteoliposome system.

We measured the uptake of 10µM [^3^H]-E_1_S by OATP1B3 proteoliposomes in the presence of 100µM EGCG. The addition of the flavonoid induced a striking increase of the initial transport rate (V_0_), causing the proteoliposomes to reach steady-state saturation in approximately 15 minutes, compared to 30 minutes for the control containing only DMSO (**Fig. 4A**). Crucially, while EGCG significantly altered the transport kinetics, it did not alter the thermodynamic steady-state, as both conditions eventually plateaued at the same luminal [^3^H]-E_1_S concentration. To determine if this direct modulatory effect was substrate-specific, we evaluated the impact of 100 µM EGCG on the transport of ^3^H-E_2_-17G. In stark contrast to the stimulation observed for E_1_S, the presence of EGCG reduced ^3^H-E_2_-17G transport to near-baseline levels (**Fig. 4B**).

**Figure 4.**
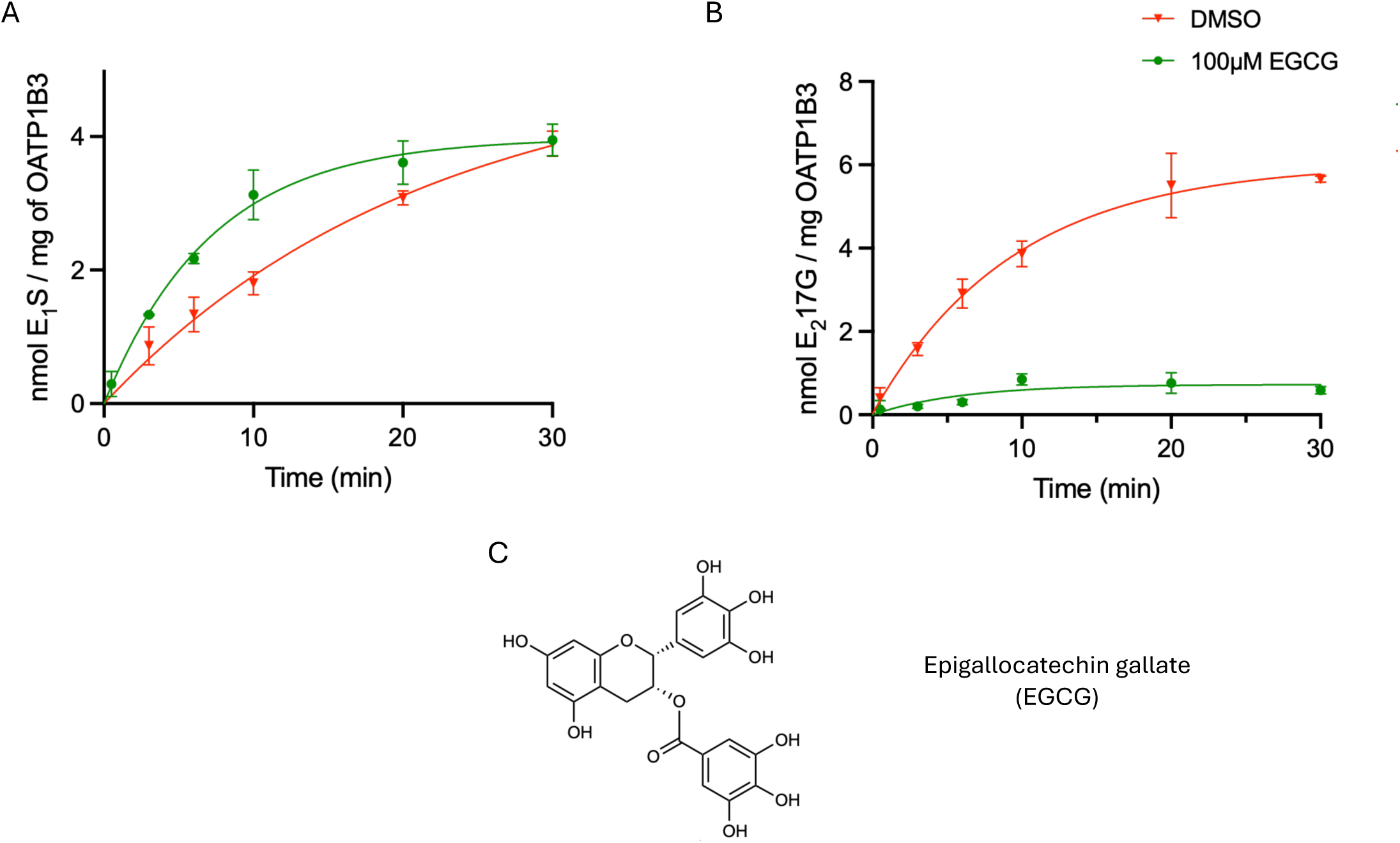
EGCG-mediated OATP1B3 uptake stimulation is substrate-specific. **A** Time-course of ^3^H-E_1_S accumulation into OTP1B3 proteoliposomes. Transport reaction were carried in the presence of 100µM EGCG or DMSO (vehicle). **B:** OATP1B3-mediated uptake of ^3^H-E_2_-17G in the presence or absence of EGCG. Data represent mean ± s.e.m. of 3 independent experiments. **C:** molecular structures of the green tea flavonoid EGCG.

## DISCUSSION

Until now, the energetic basis of OATP1B-mediated transport has remained a subject of debate, with conflicting evidence suggesting mechanisms ranging from facilitated diffusion to exchange with glutathione or bicarbonate (*10, 20, 24, 36*). This lack of consensus stems largely from the complexity of cellular experimental systems, where endogenous transport activity, intracellular metabolism, and the presence of diverse membrane proteins confound kinetic interpretations. By utilising a defined proteoliposome system, we have effectively stripped away these biological variables to interrogate the transporter in its most reductionist form.

Our findings provide direct evidence that OATP1B1 and OATP1B3 function as HCO_3_^-^exchangers in the presence of this anion. Notably, the apparent initial transport rate was largely independent of the imposed gradient, consistent with the notion that an intrinsic conformational step of the transport cycle is rate-limiting under these conditions. In contrast, steady-state accumulation was dependent on the imposed bicarbonate gradient. This distinction suggests that the bicarbonate gradient may contribute primarily to the thermodynamic bias required for uphill substrate accumulation rather than simply increasing transport velocity. We cannot exclude basal transport in the absence of HCO_3_^-^, because establishing a completely HCO_3_^-^-free environment is technically difficult owing to continuous equilibration with atmospheric CO_2_. Importantly, however, such HCO_3_^-^-free condition is not physiologically relevant. OATP1B1 and OATP1B3 are exposed to approximately 10-15mM HCO_3_^-^from the cytoplasm of the hepatocyte (*37*), indicating that bicarbonate is readily available under physiological conditions and supporting the notion that these transporters function as bicarbonate exchanger also *in vivo*. While the magnitude of this bicarbonate-driven accumulation in our isolated protein system is modest (approximately 1.4-fold), it may be interpreted not primarily as evidence for a large equilibrium gradient, but as an indication that bicarbonate coupling may help impose a net unidirectional bias on substrate uptake under physiological conditions. It is critical to interpret these values within the physiological context of the hepatocyte. In an isolated, closed proteoliposome system, transport inevitably reaches an equilibrium. *In vivo*, however, hepatocytes function as open systems in which internalized xenobiotics are continuously removed from the free intracellular pool by downstream metabolism and biliary export. Under these conditions, OATP-mediated uptake need not by itself sustain a large cytosolic substrate gradient; instead, bicarbonate coupling can help maintain net inward flux by biasing transport toward entry into a clearance pathway. Accordingly, the steady-state accumulation measured in our closed experimental system should be interpreted cautiously, as it likely underestimates the extent to which coupling can support hepatic drug uptake under physiological sink conditions. To confirm that the observed stimulation was specifically driven by a bicarbonate-dependent mechanism rather than a general anion-induced diffusion potential, we utilised NaNO_3_ as a control salt. NO_3_^-^is a chaotropic ion with significantly higher permeability than Na^+^. Therefore, when NaNO_3_ is added to the luminal space, NO_3_^-^can theoretically diffuse (as opposed to Na^+^), and generate a transient inside-positive diffusion potential (*Δψ > 0*) (*38, 39*). We observed no stimulation of substrate uptake in these conditions. This suggests that the OATP1B1 and OATP1B3 transport cycle is primarily governed by the chemical processing of the bicarbonate ion rather than by non-specific anion diffusion potentials that would favour substrate import. The direct effect of membrane potentials on OATP1B1 and OATP1B3-mediated transport in cells remains, however, a subject of debate (*22, 24*).

Capturing the reciprocal counter-ion efflux in real time remained a challenge. For this reason, we developed a functional ABCG2 and OATP1B1 co-reconstitution, which allowed us to further amplify the effect of HCO_3_^-^gradients in OATP1B1-mediated transport, as well as to directly prove for the first time the HCO_3_^-^flux as a counterion in an OATP1B system. Due to the low millimolar sensitivity of the available HCO_3_^-^probes and the low turnover rate of OATP1B1, this would not have been possible without a system that continuously resupplies substrate.

Structural analysis indicates that the bicarbonate -bound state of OATP1B3 is sterically incompatible with simultaneous substrate binding in the central, high-affinity pocket (*10*). Based on this and our transport assay data, we hypothesised that HCO_3_^-^binding is a trigger for substrate release or for preventing substrate rebinding and the subsequent reset of the transport cycle (**Fig. 5A**). In a simple facilitated diffusion model, once substrate concentrations equilibrate across the membrane, the system reaches a steady state where the transporter is equally likely to move substrate in either direction. However, in our antiport model, after substrate binding (**state V**) and translocation (**state I**), the binding of internal HCO_3_^-^to the substrate-loaded, inward-facing transporter may facilitate the release of the cargo into the cytoplasm or prevent the unproductive rebinding of the substrate (state **II**). This would avoid the re-export of the substrate into the cytoplasm. This is supported by our OATP1B1 H115A data, which revealed that HCO_3_^-^binding is essential for driving concentrative transport, as OATP1B1 retained a basal ability as a facilitated diffusion carrier when this mechanism was uncoupled (**Fig. 5B**). We therefore propose that OATP1B1 and OATP1B3 might function as a Brownian ratchet, where the transmembrane bicarbonate gradient could rectify the stochastic conformational fluctuations of the protein –typical in facilitated diffusion carriers– into unidirectional substrate flux. This allows the transporter to harness the chemical energy of the bicarbonate gradient to drive the concentrative uptake of substrates.

**Figure 5.**
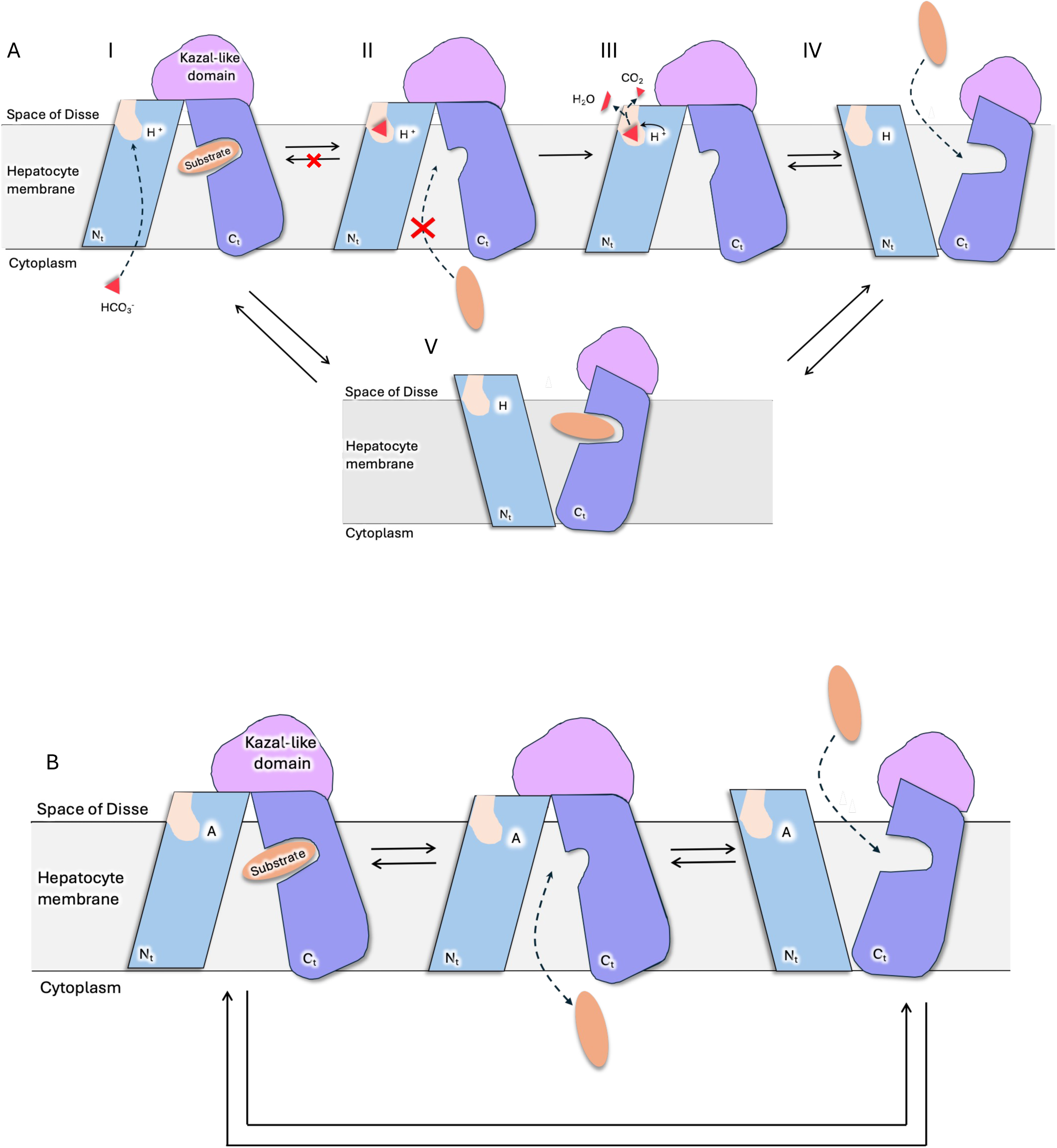
Proposed transport cycle of pH-sensitive, bicarbonate-coupled, OATP transporters. The N- and C-terminal halves, as well as the Kazal-like domain, are coloured differently. **A.** WT protein. Protonated His115 is shown as H^+^ by the bicarbonate-binding site. States I, II, IV and V have been resolved structurally (PDB 8PHW, 8PG0, 8HNB, 8HNH). We hypothesise that H115 is able to donate a proton (from the acidic space of disse) to bicarbonate, causing its hydrolysis. state III and bicarbonate hydrolysis as part of this study. **B.** H115 mutant. When the ability to coordinate bicarbonate and substrate transport is lost, we hypothesise that OATP1B1 and OATP1B3 can act merely as a diffusion facilitator.

Remarkably, the H115Q rescue variant exhibited slightly higher V_0_ than the WT while regaining sensitivity to the HCO_3_^-^gradient compared to H115A. We hypothesise that this stems from the distinct chemical properties of glutamine compared to histidine. In the WT protein, the imidazole ring of H115 likely coordinates HCO_3_ through an electrostatic interaction or salt bridge—potentially together with nearby residues such as Arg58—which effectively locks the transporter in an inward-facing conformation until the ion is processed. In contrast, the uncharged glutamine side chain in H115Q likely provides weaker hydrogen-bonding interaction. This reduced affinity might facilitate faster dissociation of the counter-ion, thereby accelerating the conformational reset and increasing the overall turnover rate in our experimental system.

This raises a fundamental evolutionary question: why would the OATP1B subfamily utilise a slower histidine-based mechanism over a kinetically superior glutamine-like one? The answer likely lies in the physiological environment of the hepatocyte. Unlike glutamine, histidine can act as a proton donor. It has been hypothesised that His115 participates in a catalytic step (*10*), facilitating the conversion of HCO_3_^-^to CO_2_ (**Fig. 5A state III**). While this step imposes a kinetic penalty, it would create a steeper counterion gradient by converting HCO_3_^-^into a neutral gas that can rapidly diffuse away to the bloodstream. Physiologically, the transmembrane HCO_3_^-^gradient in the liver is relatively shallow, from 10-15mM in the hepatocyte to lower values in the Space of Disse: due to its unstirred water layer and proton diffusion barrier nature, the pH is slightly acidic and has been estimated to be around 6.2 (*40*), and therefore HCO_3_^-^might be locally depleted (*37, 40, 41*). Thus, by catalysing the conversion to CO_2_, the WT protein would ensure that the effective concentration of HCO_3_^-^at the release site remains near zero. This maximises the thermodynamic driving force, and reduces the likelihood of reversion, allowing OATP1B1 to clear drugs from the blood even against concentration gradients—a feature that is not necessary in our proteoliposome assays, where the HCO_3_^-^experimental gradients are artificially steep and favour the non-catalytic mechanism of the H115Q mutant. This catalysis mechanism involving a histidine residue is analogous to what is seen in carbonic anhydrase (*42*) where H64 plays a key role in donating or sequestering a proton in the CO_2_ to HCO_3_^-^process and vice versa. Indeed, it has been reported that the association of carbonic anhydrase enhances the activity of other HCO_3_^-^-coupled transporters, an association known as transport metabolon (*43-46*).

This model is further supported by the natural sequence divergence within the OATP family: OATP1C1 naturally possesses a glutamine at the position corresponding to His115. Notably, OATP1C1-mediated transport is independent of pH (*20*), reinforcing the theory that a histidine residue at this position is indispensable for pH-sensitive ion coupling. Unlike the amide side chain of glutamine, the imidazole ring of histidine has a pK_a_ near the physiological range in proteins (6.9). This explains the specific pH-sensitivity of the OATP1B subfamily: more acidic extracellular environment would favour the protonated state of H115, thereby driving the catalytic cycle. This pH-dependence of the transport activity has been extensively studied in cell-based assays, where it was found that a H115A mutation abolished the pH sensitivity of OATP1B3, while the transporter did not fully lose transport activity (*20, 35, 47*). Comparable uncoupling phenotypes have been described in other transport systems. In the lactose permease LacY, mutations at Glu325 or Arg302 abolish H^+^/lactose symport while retaining exchange and counterflow, making LacY a particularly close mechanistic precedent (*48*). The multidrug transporter LmrP provides a second useful analogy: energy-uncoupled mutants show increased ethidium influx and increased drug susceptibility, consistent with uncoupling drug efflux from proton influx and permitting membrane-potential-driven drug entry (*49*). Similar separation of coupling from substrate movement has also been reported in Cl-/H+ exchangers, where uncoupling can alter the relationship between ion exchange and turnover without eliminating transport altogether (*50*). These precedents show that disrupting a proton-or ion-coupling residue does not necessarily abolish transport altogether but can instead shift the transporter toward an uncoupled transport regime.

Lastly, our findings validate our proteoliposome platform as a powerful tool for dissecting complex allosteric regulation in OATPs. While our results confirm the direct, substrate-specific stimulation of E_1_S uptake by EGCG, our observation of robust E_2_17G inhibition challenges previous cell-based reports (*26*). This discrepancy is likely explained by the significantly higher concentration of E_2_17G (30 µM) used in our assays compared to the 0.1 µM used in literature (*26*). Based on this difference, we propose two mechanistic models suggests two possible mechanistic models: (I), uncompetitive Inhibition: EGCG may act as an uncompetitive inhibitor that preferentially binds to the transporter-substrate complex. At the high substrate occupancy in our assays, EGCG would be more likely to trap the transporter, whereas at 0.1 µM, the lack of complexes would render the inhibition undetectable.

(II) Multiple Binding Sites: at high substrate concentrations, E_2_17G may occupy secondary, low-affinity binding sites within the large OATP1B3 translocation pore—sites for which EGCG might directly compete. Understanding the structural determinants that allow a single molecule to act as both an activator and an inhibitor depending on the cargo could be instrumental in the future design of drugs with tuneable OATP1B1 and OATP1B3 affinities and the prediction of complex clinical drug-drug interactions.

## METHODS

### Cell Culture and Protein Expression

Human OATP1B1 and OATP1B3 were expressed and purified as previously described (*10*). Briefly, tetracycline-inducible stable Flp-In™ T-REx™ 293 cells (Thermo Fisher Scientific) harboring C-terminally 1D4-tagged OATP1B1 or OATP1B3 were maintained in Dulbecco’s Modified Eagle Medium (DMEM, Gibco) supplemented with 10% fetal bovine serum (FBS), 100 g/mL streptomycin, and 100 units/mL penicillin at 37°C under 5%. For induction, cells were washed with phosphate-buffered saline (PBS) and transitioned to phenol red-free DMEM supplemented with 2% FBS, 1x GlutaMAX, and 1 g/mL tetracycline (Sigma). After 72 h, cells were harvested, washed in PBS, flash-frozen in liquid nitrogen, and stored at -80°C.

### Purification of OATP1B1 and OATP1B3

Cell pellets were thawed and lysed via 25 strokes in a “type B” Dounce homogenizer in lysis buffer (50 mM HEPES pH 7.4, 150 mM NaCl, 20% v/v glycerol, 1 mM PMSF, 2 g/mL DNase I, and protease inhibitor cocktail). All subsequent steps were performed at 4°C. Membranes were solubilised using 1% (w/v) n-dodecyl--D-maltopyranoside (DDM) and 0.2% (w/v) cholesteryl hemisuccinate (CHS) for 90 min, followed by ultracentrifugation at 140,000 x *g* for 45 min. The supernatant was incubated with Sepharose-coupled Rho-1D4 antibody (University of British Columbia) for 2 h. The resin was washed with 40 column volumes (CV) of purification buffer (50 mM HEPES pH 7.4, 150 mM NaCl, 20% glycerol, 0.025% DDM, 0.005% CHS). Target proteins were eluted via on-column cleavage using 3C protease (1:50 wt/wt ratio) over night.

### Purification of ABCG2

FLAG-tagged (*51*) human ABCG2 was expressed in suspension HEK293-EBNA cells via transient transfection. Cells were grown in EX-CELL® media supplemented with 2% FBS and 3 mM L-glutamine. Transfection was performed using linear polyethyleneimine (PEI) and plasmid DNA in uncomplemented FreeStyle™ 293 medium. Cells were harvested 48–72 h post-transfection. Pellets were resuspended in lysis buffer (40 mM HEPES pH 7.5, 200 mM NaCl, 10% glycerol) and solubilised with 1% DDM/0.1% CHS. After ultracentrifugation at 100,000×g, the supernatant was incubated with ANTI-FLAG® M2 affinity resin (Sigma) for 3 h. Protein was eluted using 200 μg/mL FLAG peptide and further purified via size-exclusion chromatography (SEC) on a Superdex 200 16/600 column (*52*). Peak fractions were pooled, concentrated (100 kDa MWCO), and flash-frozen.

### Proteoliposome Reconstitution

OATPs were reconstituted into a lipid mixture of PE:POPG:PC:Cholesterol (3:1:1:1 wt/wt). Lipid films were prepared via rotary evaporation, rehydrated in HBS (50 mM HEPES, 150 mM NaCl) to 20 mg/mL in a sonication bath, and subjected to three freeze-thaw cycles. Liposomes were extruded through 0.4 µm polycarbonate filters (21 times) and destabilised with 0.3% Triton X-100 (0.25% for ABCG2 co-reconstitution) (v/v). Purified protein was added at a 1:100 protein-to-lipid ratio (1:50 for co-reconstitution). Detergent was removed via sequential additions of Bio-Beads SM-2 (40 mg/mL at 15 min, 1 h, 3 h, and 80 mg/mL for 2 more hours the next day) at 4°C. Proteoliposomes were harvested by centrifugation (280,000 x *<u>g</u>*, 20 min) and resuspended in HBS. Reconstitution efficiency was assessed using the Schaffner and Weissmann assay (*53*), ensuring that WT and mutant data would be normalised to protein concentration.

### In vitro Transport Assays

#### Radiolabelled Assays

Proteoliposomes underwent five freeze-thaw cycles in the internal buffer, were extruded (0.4 μm, 21 times), and pelleted (100,000×*g,* 30 min). They were resuspended in outside buffer. Uptake was initiated by diluting proteoliposomes (4 mg/mL) into pre-warmed HBS containing 30 μM ^3^H-E_2_17G (OATP1B3) or 10 μM ^3^H-E_1_S (OATP1B1). At specified intervals, 50 μL aliquots were quenched in 600 μL ice-cold HBS containing 80 μM unlabelled substrate, filtered through 0.22 μm PVDF membranes, and washed twice. Radioactivity was quantified via liquid scintillation counting. OATP1B1 transport assays were carried at 25°C and for OATP1B3, 30°C.

#### HPLC-based Statin uptake Assays

Transport of 100 μM fluvastatin was performed at 27°C for 20 min. Quenched samples were filtered and extracted in a 1:1 acetonitrile/methanol mixture. Fluvastatin was quantified via reverse-phase HPLC (C18 column, 1.5 μm, 100 x 2.1 mm) with UV detection at 240 nm. A gradient of (A) 20 mM ammonium acetate (pH 4) and (B) acetonitrile, 0.1% acetic acid) was employed. The run consisted of a gradient with a gradient of phase B from 40% at 0min to 70% at 3.9min. From 3.9 to 5 min the column was re-equilibrated with 40% phase B (Flow rate 0.6mL/min). A standard curve was made using pure fluvastatin in DMSO

#### Co-reconstitution and Luminescence Assays

For ABCG2-OATP1B1 relay assays, the external buffer was supplemented with 5 mM ATP and 5 mM MgCl_2_. At each timepoint the sample was filtered in a 96-well 0.65-1µM glass fibre filter plate and washed twice with 200µL.

In luminescence-based assays, proteoliposomes were loaded with 50 μM [Eu.L1]^+^ probe and 100 μM E_1_S. Liposomes were pelleted and resuspended in HBS supplemented with 100 µM E_1_S. The final assay buffer consisted of HBS supplemented with 15mM NaHCO_3_, with 5mM ATP, 5mM MgCl_2_, 2U/mL creatine phosphokinase, 50µM creatine phosphate and 50µM E_1_S. The assay was carried in a black, transparent bottom 96-well plate at 37°C. At each time point, Eu(III) luminescence was measured in a Synergy Plate Reader using (λ_ex_ =332 nm, λ_em_=617 nm) (time-resolved emission using a 100 μs lag, and 500 μs collection) at 37°C. Luminescence intensity was normalised to the average intensity at timepoint 5 min, and then it was corrected by the relative luminescence loss measured in a liposome-only control under the same experimental conditions.

### Liposome luminal volume estimation

The luminal volume of the liposomes was calculated following a previously described protocol (*54*). The liposomes in aqueous buffer where freeze dried and resolubilised in diethyl ether. This solution was dried in a Rotary evaporator, and the fil was rehydrated in HBS with 0.5µM calcein in a sonication bath. The liposomes were freeze-thawed 5 times and extruded 21x through a 0.4 µm filter. The liposomes were diluted to 0.5mg/mL of lipids and the florescence was measuresd (λ_ex_= 490 nm, λ_em_= 520 nm) in a transparent bottom, black 96-well plate. Then the florescence of the luminal calcein outside the liposomes was quenched with 10mM CoCl_2_ and luminal florescence was measured again. To measure the base line, 1.5% Triton-X 100 was added to solubilise the liposomes and florescence was measured again. The ratio of internal vs. total fluorescence was used to calculate a luminal volume of 0.85 ± 0.02 µl mg^-1^ of lipids.

### Statistical Comparisons and figure preparation

Analyses were performed in GraphPad Prism v10 (GraphPad Software). When appropriate, data was fit to one-phase association equations Y=Y_0_ + (Plateau-Y_0_) x (1-e^(-K*X)^). Graphs were made in GraphPad Prism 10.1.1 and structural figures in ChimeraX (*55*).

## AUTHORS CONTRIBUTION

K.P.L. and J.C.M.T conceived the study. J.C.M.T performed the experiments: protein expression and purification, reconstitution and transport assays. S.J.B. Developed the lanthanide probe. K.P.L, B.S. and J.C.M.T wrote the manuscript with input from all authors.

## ETHICS DECLARATION

The authors declare no competing interests.

## ACKNOWLEDGEMENTS

We thank Raimund Dutzler for helpful discussions. This work was supported by the Swiss National Science Foundation (grant 214834 to K.P.L).

