## Supplemental Figures 1-2 for "Functional Reconstitution of Human OATP1B1 and OATP1B3 in Proteoliposomes: Effect of Bicarbonate Gradients and Substrate-specific Stimulation on Transport"

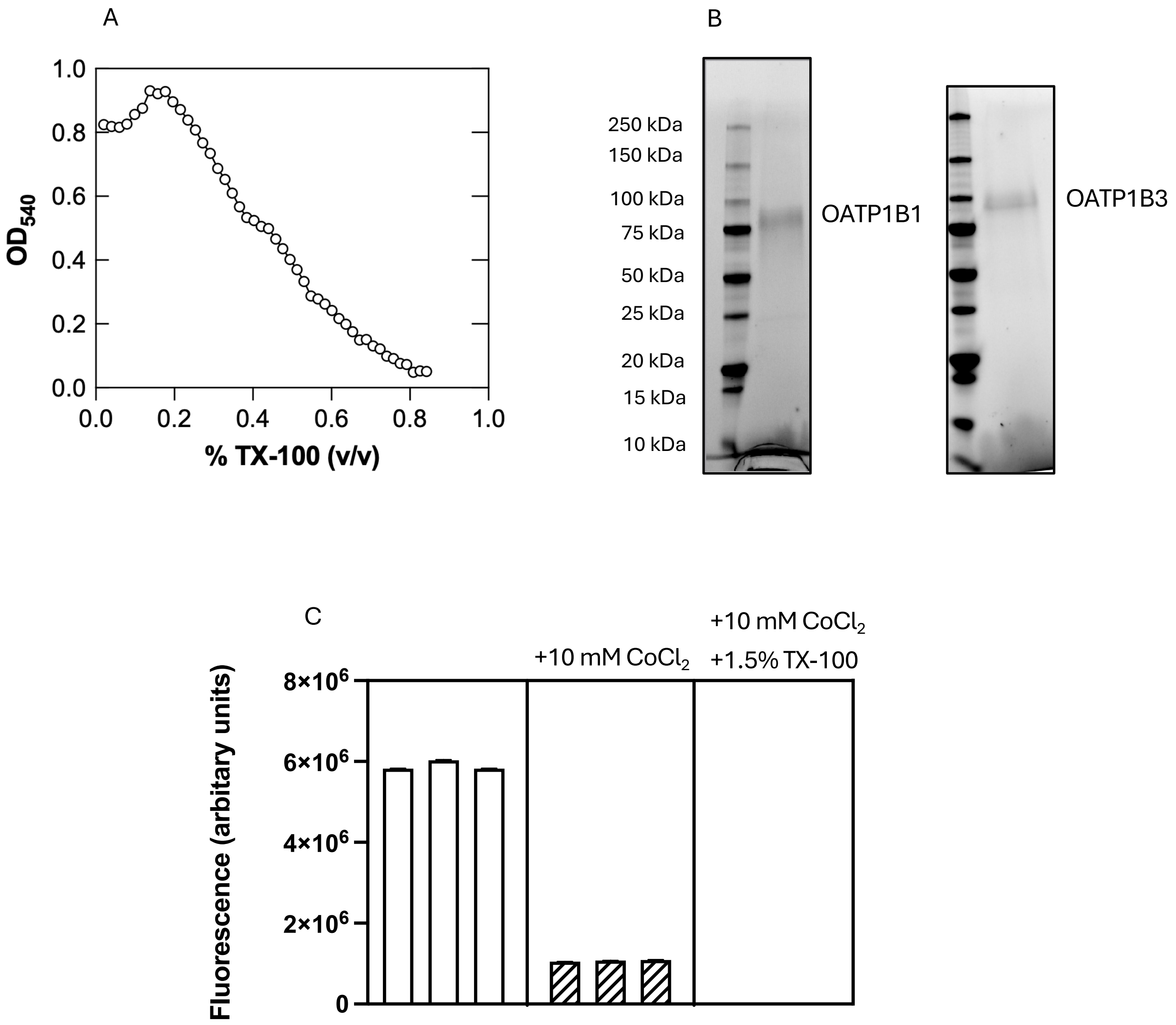


**Supplementary Figure 1**. **Liposome reconstitution and determination of the liposome luminal volume.** **A.** Liposomes destabilisation profile, to estimate the concentration of Triton X-100 required for reconsititution. **B.** SDS-PAGE gel of the proteoliposomes containing OATP1B3 (right) and OATP1B1 (left). **C.** Calcein fluorescent signals of the proteoliposomes solution before quenching with $CoCl_2$, after quenching and after solubilising the liposomes. Each bar respresents a biological replicate, each measured in six technical replicates.

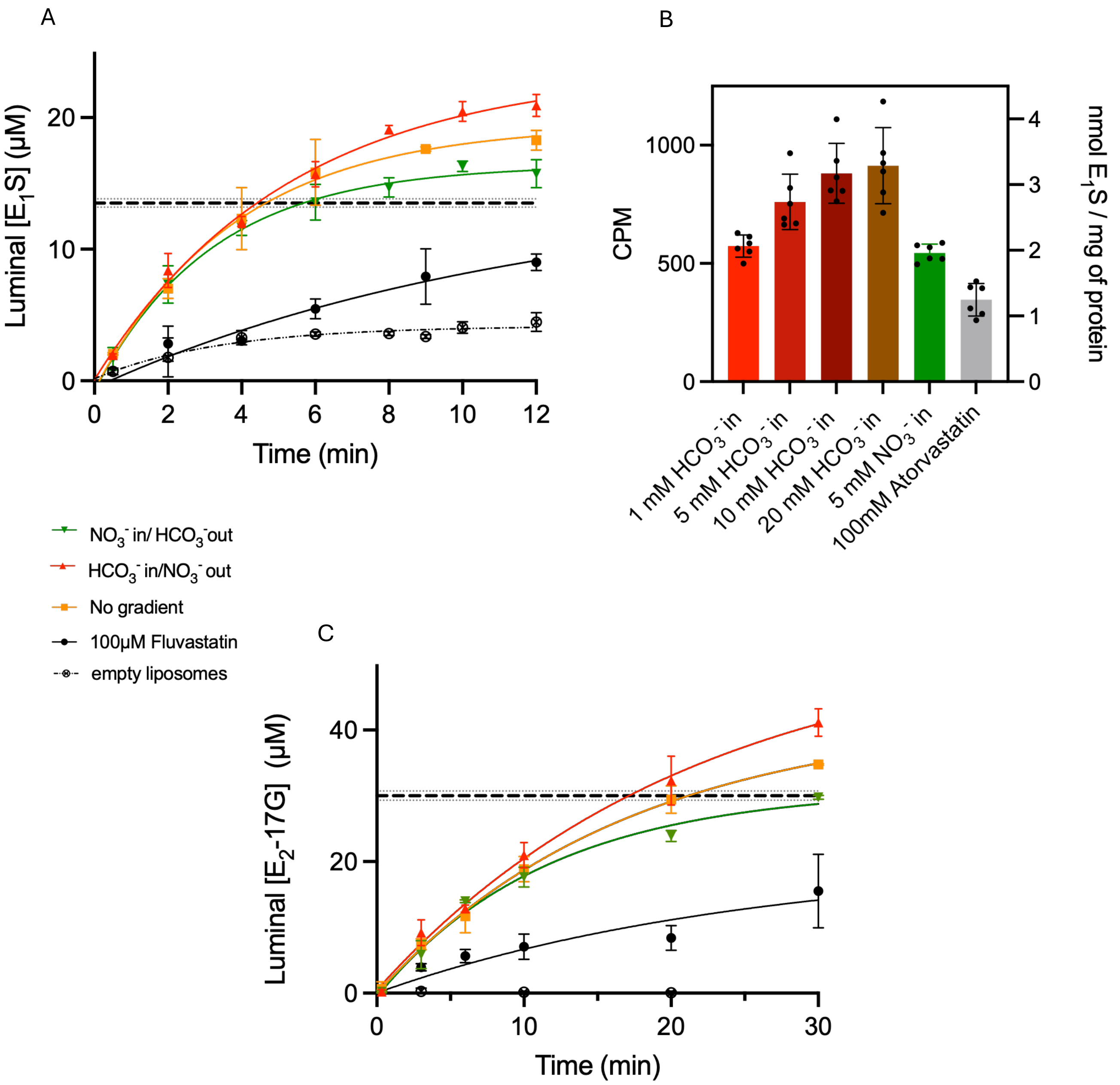


**Supplementary Figure 2**. **Uptake of Radioactively labelled substrate by OATP1B1 and OATP1B3 A.** Time-course of $^{3}$H-$E_1$S uptake by OATP1B1, including protein-free liposomes as control for unspecific substrate adsorption. **B.** $^{3}$H-$E_1$S uptake by OATP1B1 liposomes in response to different magnitudes of $HCO_3^-$ gradient, after 15min. **C.** Time-course of $^{3}$H-$E_2$17G uptake by OATP1B3, including protein-free liposomes as control for unspecific substrate adsorption.
